# Genomic Skimming and Nanopore Sequencing Uncover Cryptic Hybridization in One of World’s Most Threatened Primates

**DOI:** 10.1101/2021.04.16.440058

**Authors:** Joanna Malukiewicz, Reed A Cartwright, Jorge A Dergam, Claudia S Igayara, Patricia A Nicola, Luiz MC Pereira, Carlos R Ruiz-Miranda, Anne C Stone, Daniel L Silva, Fernanda de Fátima Rodrigues da Silva, Arvind Varsani, Lutz Walter, Melissa A Wilson, Dietmar Zinner, Christian Roos

## Abstract

The Brazilian buffy-tufted-ear marmoset (*Callithrix aurita*), one of the world’s most endangered primates, is threatened by anthropogenic hybridization with exotic, invasive marmoset species. As there are few genetic data available for *C. aurita*, we developed a PCR-free protocol with minimal technical requirements to rapidly generate genomic data with genomic skimming and portable nanopore sequencing. With this direct DNA sequencing approach, we successfully determined the complete mitogenome of a marmoset that we initially identified as *C. aurita*. The obtained nanopore-assembled sequence was highly concordant with a Sanger sequenced version of the same mitogenome. Phylogenetic analyses unexpectedly revealed that our specimen was a cryptic hybrid, with a *C. aurita* phenotype and *C. penicillata* mitogenome lineage. We also used publicly available mitogenome data to determine diversity estimates for *C. aurita* and three other marmoset species. Mitogenomics holds great potential to address deficiencies in genomic data for endangered, non-model species such as *C. aurita*. However, we discuss why mitogenomic approaches should be used in conjunction with other data for marmoset species identification. Finally, we discuss the utility and implications of our results and genomic skimming/nanopore approach for conservation and evolutionary studies of *C. aurita* and other marmosets.

## Introduction

‘Genomic skimming’ is an emerging technique involving random shotgun sequencing of a small percentage of total genomic DNA, which results in a comparatively large sampling of high-copy portions of the genome like the mitogenome and repetitive elements^1^. As genetic characterization of a species is fundamental for conservation and evolutionary studies, this technique is expanding the use of mitogenomics for phylogenetics, species identification, and biodiversity assessments of under-studied organisms^2,3^. While genomic skimming originates from short-read, next generation sequencing, recent implementations of the technique instead have utilized portable, long-read nanopore sequencing to successful generate *de novo* mitogenomic assemblies of genomic-data deficient vertebrates without the need for PCR^4, 5^. Thus, the coupling of genomic skimming with nanopore sequencing holds great potential to address genomic data deficiencies in non-model organisms to assist in species conservation efforts and evolutionary studies.

In this work, we aimed to develop a rapid, cost-effective protocol to obtain genomic data for one of the world’s most endangered primates, the small buffy-tufted-ear marmoset *C. aurita* (Figure 1), for which there is relatively little genetic knowledge. This species is native to the mountainous regions of southeastern Brazil^6,7,8^, and there are about 10,000 mature *C. aurita* remaining in the wild^9^. The IUCN Red List considers this species Endangered, and *C. aurita* conservation threats include habitat deforestation, urbanization, and logging^9^. The buffy-tufted-ear marmoset also faces ecological and hybridization threats from two invasive marmoset species that have been introduced by humans into southeastern Brazil, *C. jacchus* and *C. penicillata* (natural ranges and species shown in Figure 1)^7,8,9,10,11^. Currently, a captive breeding program is underway for *C. aurita*, with the aim to eventually reintroduce the species into the wild. As genetics and genomics are an integral part of the *C. aurita* program for species identification, breeding, and population management^8, 12^, rapid access to genetic data is crucial for the long-term success of *C. aurita* conservation.

**Figure 1.**
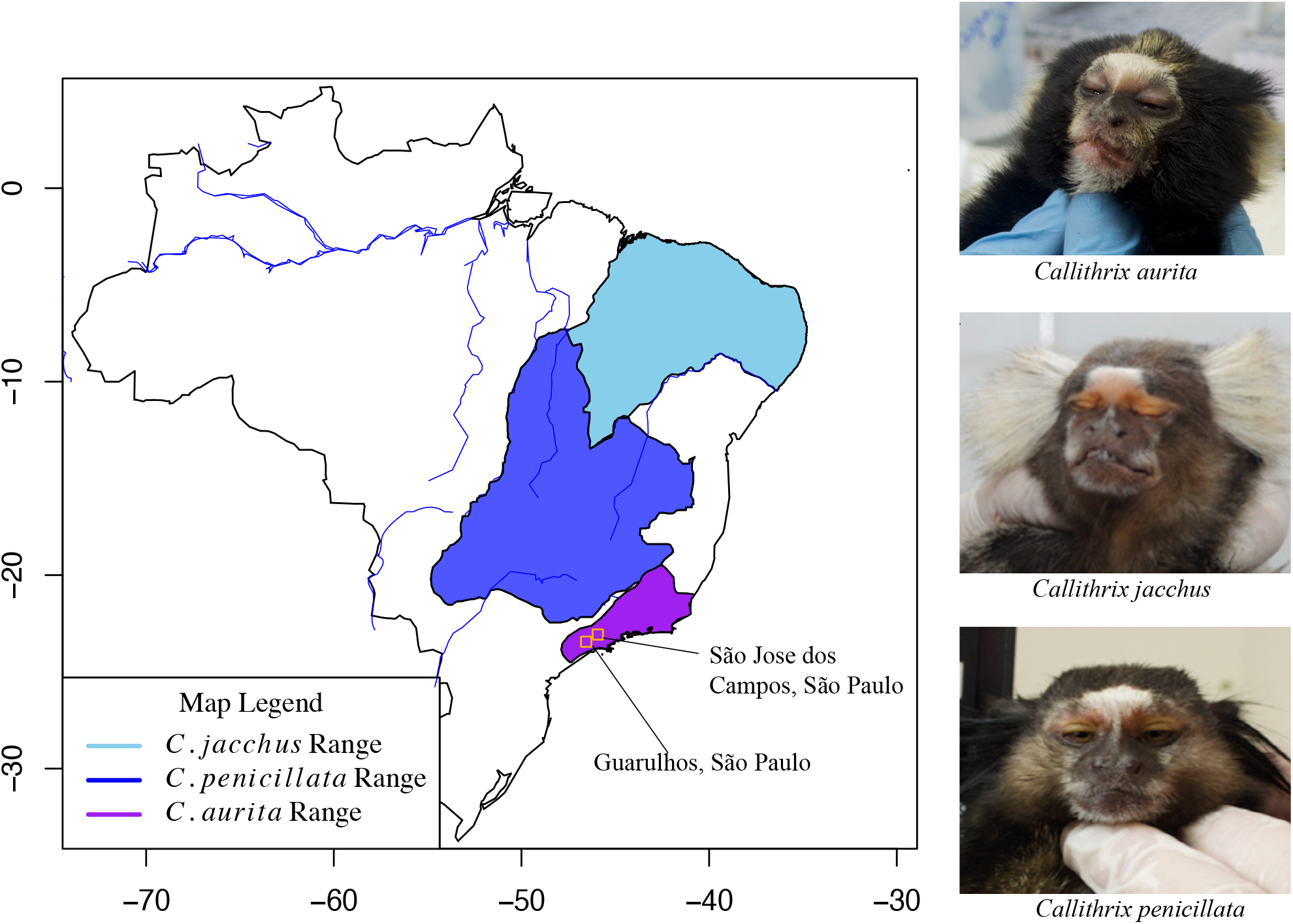
The distinct natural distributions and phenotypes of *C. jacchus*, *C. penicillata*, and *C. aurita*. Cities of origin (São Jose dos Campos) and collection (Guarulhos) of individual BJT022 are shown as orange squares. The photographs on the right hand side show the phenotype of individual BJT022 as matching that of *C. aurita*, as well as the phenotypes of *C. jacchus* and *C. penicillata* for reference.

Most *Callithrix* genetic work is based on phylogenetic analysis of short mitochondrial DNA (mtDNA) regions such as *COI*, *COII*, and the control region to resolve phylogenetic relationships (e.g.,^10,13,14,15^). However, due to the relatively recent divergence times between *Callithrix* mtDNA lineages^8,16^, these *Callithrix* phylogenies usually show polytomies or lack strong statistical support. Nonetheless, mitogenomics alleviates some of these difficulties in *Callithrix* phylogenetics studies^7,16^, and the *C. aurita* mitogenome was recently assembled and annotated with Sanger sequencing and short-read high-throughput sequencing^7^. However, one drawback to current *Callithrix* mitogenomics work is the use of relatively time-consuming molecular techniques and lag time for sequencing data generation from several weeks to months (pers. obs., Malukiewicz).

As a subsequent step in developing genomic resources for *C. aurita*, we present here a rapid, PCR-free, genomic skimming technique for mitogenomic reconstruction with the hand-held, long-read Oxford Nanopore Technologies (ONT) minION sequencer. We assessed whether the mitogenome of a single buffy-tufted-ear marmoset could reliably be reconstructed with ONT sequencing, and then be used for downstream phylogenetic and genetic diversity analyses. We checked the accuracy of the ONT mitogenome assembly against a mitogenome assembly generated from the same marmoset individual using long range PCR and Sanger sequencing^7,16^. There was not only very high concordance between the ONT and Sanger-derived sequences, but with this technique we also uncovered unexpected discordance between the phenotype and mitogenomic lineage of the sampled marmoset individual. Our analyses showed that our marmoset individual was actually a cryptic hybrid possessing a mitogenome lineage belonging to *C. penicillata*, but a *C. aurita* phenotype. This finding represents the first recorded case of genetic introgression of *C. penicillata* genetic material into *C. aurita*. Our approach involving ONT-based genomic skimming increased the available genomic information for *Callithrix* taxa and provides a robust and accessible tool for genomic studies and conservation of *Callithrix*, particularly *C. aurita*.

## Results

### Phenotypic Taxon Identification of Sampled Marmoset Individual

We collected a small skin biopsy from a single individual (BJT022) housed at the Guarulhos Municipal Zoo, Guarulhos, São Paulo state, weighed the individual, and photographed it. Zoological records state that the individual originates from São Jose dos Campos, SP (Figure 1). Based on previous phenotypic and morphological description of the species^7,8,17^, we initially identified the individual as *C. aurita*. The individual had the yellow ear tufts characteristic of *C. aurita*, as well as yellow pelage above the forehead, and other yellow patches around the facial region^17^. The overall pelage of the individual was black, with yellow pelage prominently visible, but intermixed with darker pelage at the hands. As can be seen from Figure 1, the phenotype of *C. aurita* is distinct from that of *C. jacchus* and *C. penicillata*, which are commonly found as invasive, exotic species in the natural range of *C. aurita*. Morphologically, the individual was within the expected body weight range for *C. aurita* (approximately 450g)^8^. Other *Callithrix* species tend to weight relatively much less (200-300g)^8^.

### Complete Mitogenome Reconstruction

After direct ONT sequencing of genomic DNA (gDNA) from individual BJT022, we successfully determined a full mitogenome using the genomic skimming/minION approach. ONT sequence reads were first filtered for quality and to remove sequencing adapters. Then ONT mtDNA reads were identified with the BLAST-based mtBlaster script^5^, with the *C. aurita* (GenBank Accession MN787075) mitogenome as the reference genome. Table 1 shows overall characteristics for gDNA ONT reads as well as ONT mtDNA reads. The final length of the reconstructed ONT-derived mitogenome was 16,181 base pairs (bps) with an average coverage of 9x. Annotation of the ONT-derived mitogenome with the CHLOROBOX web server^18^ showed that the mitogenome contained 37 mitochondrial genes (13 protein-coding genes, 22 tRNAs, and two rRNAs), as shown in Figure 2.

**Table 1.** Metrics for BJT022 minION mitogenome assembly

| Information | Result |
| --- | --- |
| Total Number of Genomic Reads | 50553 |
| Mean Genomic Read Length | 3208.31 |
| Mean Genomic Read Quality | 12.2 |
| Average Read Length N50 | 4669.63 |
| Number of mtDNA Reads | 60 |
| Average mtDNA Read Length (bp) | 3866.75 |
| Average mtDNA Base Quality (Phred score) | 20.44 |
| Average mtDNA Sequencing Depth | 9x |
| Final mtDNA Genome Length (bp) | 16181 |

**Table 2.** *Callithrix* Diversity Measures

| Species | Sequences | Haplotypes | Haplotype Diversity | Nucleotide Diversity | Theta | Variable Sites |
| --- | --- | --- | --- | --- | --- | --- |
| *C. aurita* | 6 | 4 | 0.87 | 0.00315 | 44.67 | 102 |
| *C. geoffroyi* | 7 | 4 | 0.81 | 0.00535 | 78.37 | 190 |
| *C. jacchus* | 9 | 7 | 0.94 | 0.00109 | 23.92 | 65 |
| *C. penicillata* | 8 | 7 | 0.96 | 0.01166 | 162.75 | 414 |

**Figure 2.**
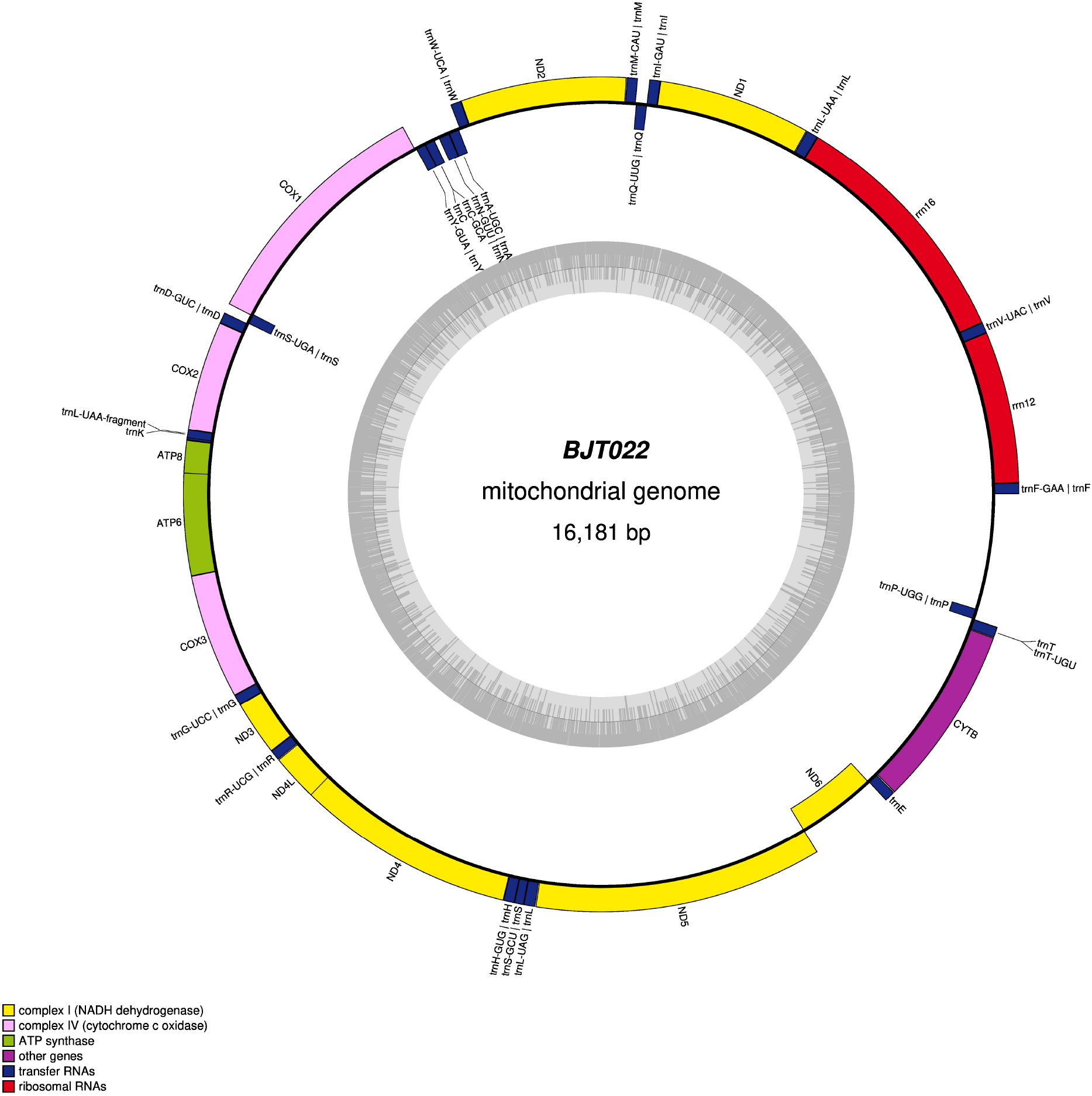
Circular map of the *Callithrix* mitogenome obtained from ONT sequencing. Color legend in bottom left corner indicates annotated features of the mitogenome assembly, and the inner circle within the plot shows sequence %GC content.

We assessed accuracy of the reconstructed ONT-derived mitogenome by comparing the resulting sequence with that of a Sanger-sequenced mitogenome from the same individual. To note, the Sanger sequence mitogenome was missing 119 bases that comprised the entire tRNA-Phe and part of the 12S rRNA region. Nonetheless, pairwise-alignment of the Sanger and ONT-derived sequences of the BJT022 mitogenome showed 99.4% concordance, as illustrated in the dot plot in Figure 3. Sites of non-concordant bases between the two sequences accounted for 0.002% of overall alignment sites. Then sites with missing bases, mostly within long homopolymer runs within the ONT sequence, accounted for 0.004% of all alignment sites. To finalize the mitogenome assembly of BJT022, we made a consensus sequence by using the Sanger sequence to correct for non-concordant sites and homopolymer runs, and the ONT sequence to fill the 119 bases missing from the Sanger sequence. This consensus alignment had a nucleotide composition of A 32.6%, T 27.0%, C 27.0%, and G 13.4%.

**Figure 3.**
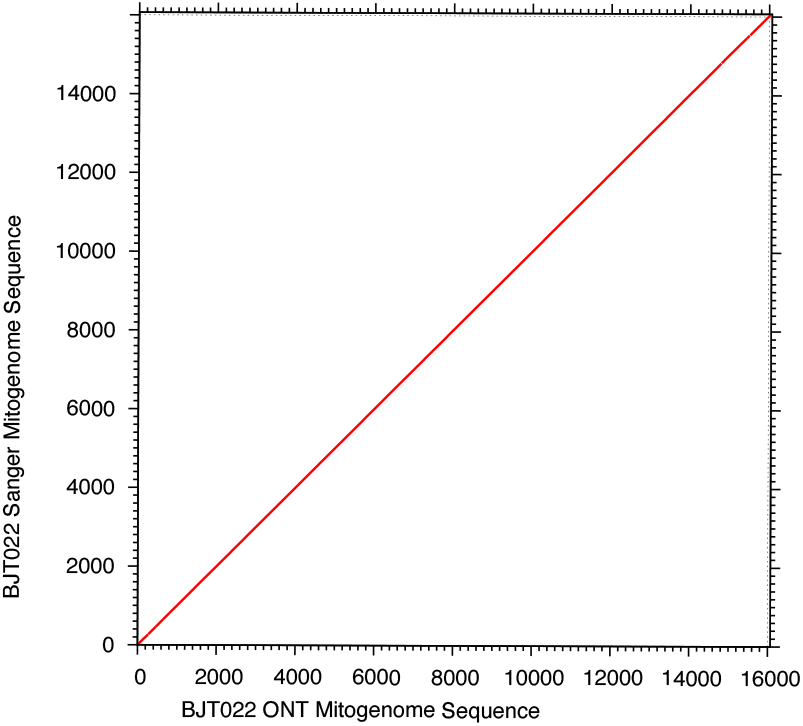
Dot plot of pairwise alignment of the mionION and Sanger sequence *Callithrix* mitogenome of marmoset BJT022. The alignment was produced in MAFFT.

### Phylogenetic Reconstruction

We obtained a well supported Maximum Likelihood (ML) phylogeny (Figure 4, Figure S1) that combined the new BJT022 consensus mitogenome sequence with previously available *Callithrix* and other New World primate mitogenome sequences^7^. As we obtained a phylogeny with a highly similar topology and levels of statistical support to that of Malukiewicz et al. (2021)^7^, we used the same phylogenetic clade designations for *Callithrix* mitogenomic haplotypes as in that previous study. In examining the *C. aurita* clade within the resulting phylogeny, two distinct geographical clusters appear, one composed of two haplotypes originating from Rio de Janeiro state, and the other cluster composed of a mix of haplotypes originating from Minas Gerais and São Paulo states, to the exclusion of a haplotype from Rio de Janeiro states (following provenance information in Malukiewicz et al. (2021)^7^).

**Figure 4.**
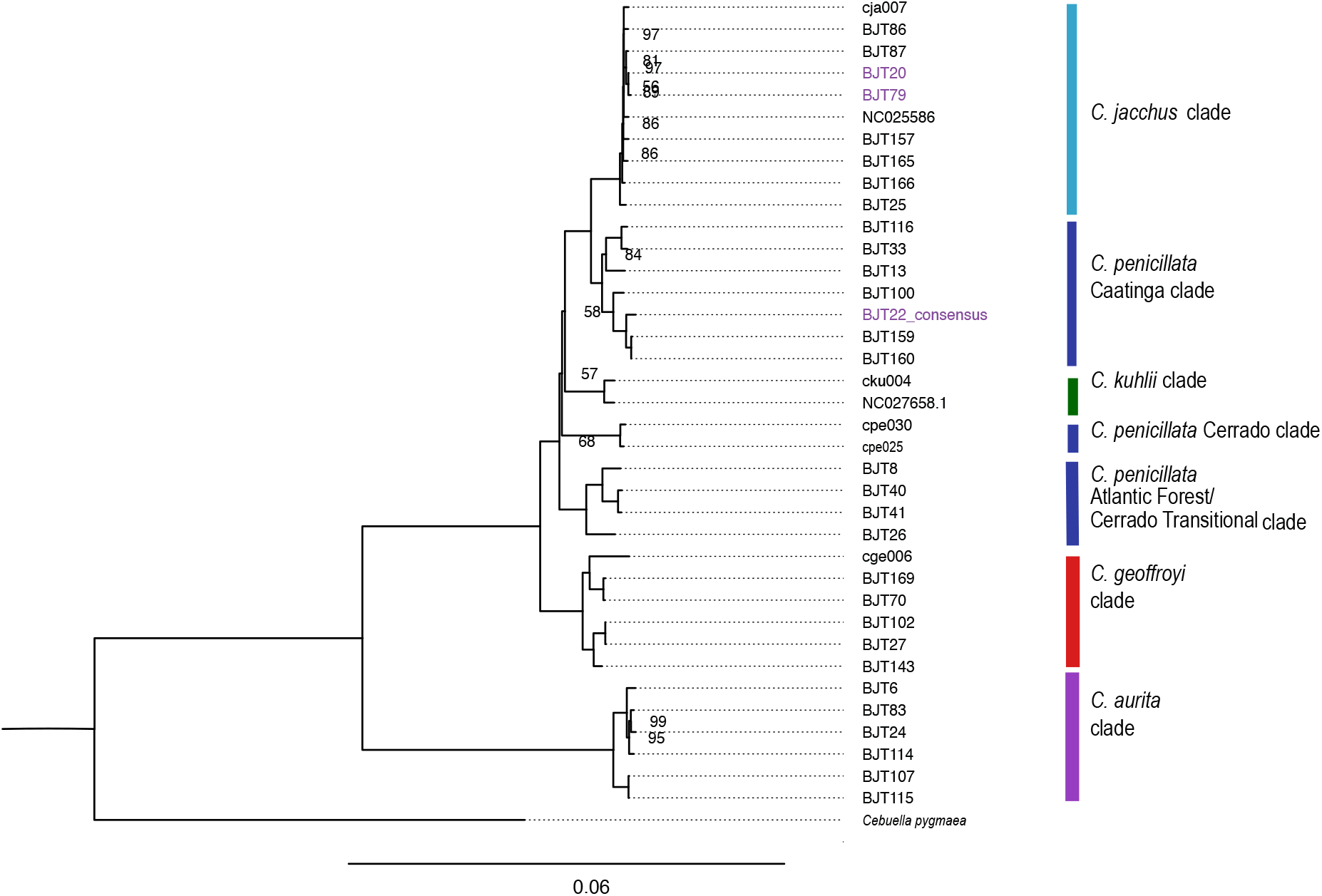
ML tree showing phylogenetic clustering of the *Callithrix* mitogenome of marmoset BJT022 within the *C. penicillata* Caatinga clade (complete tree with outgroups is presented in Figure S1). The BJT022 mitogenome is highlighted at the tree tips in purple along with two other mitogenomes of marmosets with *C. aurita* phenotypes from Malukiewicz et al. (2021)^7^ that were incongruent with their mitogenmic lineages. Categorization of major *Callithrix* mitogenomic clades follow that of Malukiewicz et al. (2021).^7^

In our phylogeny, the mitogenomic lineage of individual BJT022 unexpectedly clustered not within the *C. aurita* clade, but rather with the *C. penicillata* Caatinga clade. Additionally, two mitogenome lineages from the Malukiewicz et al. (2021)^7^ study that originated from two marmosets with *C. aurita* phenotypes clustered within the *C. jacchus* clade in the previous phylogeny as well as in ours. Thus, these two studies show three cases of cryptic marmosets hybrids with discordant *C. aurita* phenotypes and non-*C. aurita* mitogenomic lineages.

### *Callithrix* Mitochondrial Genetic Diversity

We used *Callithrix* mitogenome sequences from Malukiewicz et al. (2021)^7^ to calculate diversity indexes of four *Callithrix* species (*C. aurita*, *C. jacchus*, *C. geoffroyi*, and *C. penicillata*), for which haplotypes were available for three or more individuals per species (Table 4). Relative to the other four species, *C. penicillata* showed much higher values across all diversity indexes (Table 4). *Callithrix jacchus* showed among the highest haplotype diversity values, but then had the lowest values for all other diversity indexes. Diversity indexes for *C. aurita* generally fell in the middle relative to values of the other species. This was also the case for *C. geoffroyi* diversity indexes.

## Discussion

### Feasibility of Genomic Skimming on the ONT minION Sequencer

The term ‘genomic skimming’ was first coined by in 2012 by Straub et al. (2012)^19^ as a way to utilize shallow sequencing of gDNA to obtain relatively deeper coverage of high-copy portions of the genome, including mitogenomes. In combining genomic skimming with ONT long-read sequencing, we successfully reconstructed a complete marmoset mitogenome without the need for prior PCR enrichment, using standard molecular biology equipment, and a compact portable sequencer that connects to a laptop computer. Preparation of genetic material for ONT sequencing in this study took less than a full day, and sequencing reads were available within 48 hours. Although the coverage of our reconstructed ONT mitogenome was low-medium (9x) and one of the largest sources of error for the ONT reads were missing reads in long homopolymer runs, the ONT data showed a high degree of concordance with gold standard mtDNA Sanger sequencing reads for the same individual. Hence, this work along with a number of previous studies (e.g.,^5, 20^), highlights ONT-based genomic skimming as holding great potential for enhancing mitogenomic and diversity studies of data-deficient and/or non-model organisms.

A major challenge in ONT sequencing is the relatively high sequencing error (5%-15%), but the application of computational ‘polishing’ significantly reduces errors of raw ONT data (e.g.^21^). Another challenge with ONT methodologies is the large amount of input DNA needed for sequencing relative to other types of methods, particularly PCR and Sanger sequencing. Multiplexing samples onto the same flow cell is one way to reduce the required amount of per sample DNA, and currently ONT chemistry allows for up to 24 individual gDNA samples to be multiplex per flow cell. Another option to improve mitogenome coverage from genome skimming shotgun data, especially for sensitive applications is to use sample preparation approaches that specifically enrich for mtDNA (e.g., <https://www.protocols.io/view/isolation-of-high-quality-highly-enriched-mitochon-mycc7sw>).

It is important to point out that our approach represents a starting point from which methodological aspects could be adjusted to further improve and modify our protocol. An important consideration for long-read sequencing is access to high-quality DNA which is not degraded. For marmosets especially, another consideration for input DNA is whether chimerism could bias genomic analysis or not, as levels of chimerism vary between marmoset biological tissues. Marmosets usually give birth to twins that are natural hematopoietic chimeras due to cellular exchange from placental vascular anastomoses during early fetal development^22,23,24^. This chimerism may result in the presence of up to 4 alleles of a single-copy genomic locus within a single individual. In marmosets, skin shows some of the lowest amounts of chimerism while blood is highly chimeric^23,25,24^. Depending on project design, high levels of chimerism can bias base calling of nuclear genome derived sequence reads, but this is less of a concern for mitogenomic studies as mtDNA is haploid and transmitted maternally.

In this work, we obtained DNA from a ear skin biopsy, but this represents a minimally invasive source of genetic material. As an epidermal tissue, buccal swabs are a relatively less invasive source of low-chimerism epidermal DNA. Recently, urine has also been shown to be a non-invasive source of high-quality DNA^26^, but the amount of chimerism is currently not known for marmoset urine. Urine represents a potentially non-invasive genetic tissue which could be combined with genomic skimming of highly endangered non-model organisms, particularly within captive settings.

### *Callithrix aurita* and Anthropogenic Marmoset Hybridization

Our original aim in this work was to reconstruct the mitogenome of the endangered buffy-tufted-ear marmoset with a PCR-free ‘genomic skimming’ approach with minimal technical requirements. We successfully reconstructed the full mitogenome from a captive individual possessing a *C. aurita* phenotype, but the mitogenomic lineage showed unexpected discordance with this phenotype. While we expected the mitogenome of the sampled individual to be that of *C. aurita* instead the sampled individual possessed a *C. penicillata* Caatinga clade mitogenome lineage. These results indicate that our sampled marmoset was actually a cryptic *C. aurita* x *C. penicillata* hybrid, and represents the first ever genetic instance of observed one-way genetic introgression from *C. penicillata* into *C. aurita*. More specifically, the results show introgression of exotic female *C. penicillata* into native *C. aurita*.

Although *Callithrix* species are naturally allo- and parapatric, *C. penicillata* and *C. jacchus* have been introduced into the native range of *C. aurita* in southeastern Brazil largely as a result of the illegal pet trade and subsequent releases of exotic marmosets into forest fragments^7,8^. The *Callithrix* genus is a recent primate radiation, and while divergence times between *Callithrix* species vary, all *Callithrix* species can still interbreed^11^. Malukiewicz et al. (2021)^7^ recently found evidence of genetic introgression from of exotic *C. jacchus* into *C. aurita* within the metropolitan area of the city of São Paulo. This case was similar to the results presented here in that there was discordance between the individuals’ *C. aurita* phenotypes and the phylogenetic lineage of their mitogenome. One of these previously sampled cryptic *Callithrix* hybrids originated from the wild in the municipality of Mogi das Cruzes, which lies in the eastern portion of metropolitan São Paulo^27,8^. Following zoological records, the animal we sampled here originated from the municipality of São Jose dos Campos, which also lies in the eastern portion of metropolitan São Paulo.

The above results are alarming since they suggest that genetic introgression is underway from exotic, invasive marmosets to the endangered, native marmosets of southeastern Brazil. At this time, it is not possible to determine how board this pattern is at the geographic, genomic and species levels, and whether introgression is only unidirectional and exactly which exotic and native species are involved. Specifically for *C. aurita*, unidirectional genetic introgression from invasive marmosets as well as cryptic hybridization is worrying due to the species’ threatened conservation status. A small number of captive facilities around southeastern Brazil are currently breeding captive *C. aurita* for eventual reintroductions into the wild^12,8^. Individuals within these captive populations should be confirmed both genetically and phenotypically as not being of hybrid origin, as to avoid introducing exogenous genetic material into the captive population and subsequently into the wild. Additionally, further genetic information is need on wild *C. aurita* populations to not only characterize diversity within the species, but also to better assess the occurrence of hybridization between exotic and native marmosets in southeastern Brazil. This information is critical for defining genetic diversity of *C. aurita* and maintaining species genetic integrity in the wild and captivity.

### Utility of Mitogenomics for Evolutionary and Conservation Studies of *Callithrix aurita* and Other Marmosets

The buffy-tufted-ear marmoset is not only critically endangered but also highly data-deficient in terms of genetic information. The limited number of genetic studies involving *C. aurita* have used the mtDNA control region^13,15^, *COI*^10^, and the mitogenome^7^ for phylogenetic study of *Callithrix* mtDNA lineages, species identification, and detection of hybridization. The phylogenies obtained by us and Malukiewicz et al. (2021)^7^ do show some geographical separation between *C. aurita* mitogenome haplotypes originating from different portions of the species’ natural range. Our calculation of *Callithrix* mtDNA diversity indexes based on data from Malukiewicz et al. (2021)^7^ show that diversity in *C. aurita* is still comparable to that of other *Callithrix* species. However, a large sampling effort of *C. aurita* in terms of individual numbers and across the species range is needed for accurate determination of current levels of species standing genetic variation. Additionally, surveys should be conducted of the standing genetic variation levels of the captive *C. aurita* population. These data are crucial for understanding anthropogenic impacts on the species as well for making appropriate decisions for species conservation.

The application of genomic skimming based on portable ONT long-read technology can be applied to address several of these knowledge gaps for *C. aurita*. First, with large-scale sampling of wild and captive *C. aurita*, genetic diversity estimates, demographic history, and other evolutionary analyses can be calculated relatively easily from mitogenomic data. Given the relatively fast turnaround time to obtain sequencing data from the minION, such data could be quickly obtained for a primate as highly endangered as *C. aurita*, without weeks or months long wait times for sequencing data. Laboratory setup of the minION also does not require any additional special equipment, which also makes genomic work with highly endangered species as *C. aurita* accessible for investigators under relatively constrained budgets.

*Callithrix aurita*’s sister species *Callithrix flaviceps* faces a similar plight as *C. aurita*, but with an adult population estimated to be at about 2000 adult individuals^28^. Currently there are also plans to breed *C. flaviceps* in captivity for eventual wild reintroduction, but currently there is, to our knowledge, no genetic data available for this species. Thus, the same sort of sampling and research efforts are needed for *C. flaviceps* as for *C. aurita*, perhaps even more urgently for the former species given its smaller population. As such, *C. flaviceps* is a good candidate case for the adaptation of techniques such as genomic skimming and low-cost desktop sequencing to rapidly increase genomic resources for a non-model species for conservation and evolutionary studies.

In the case of marmosets, while mitogenomics shows great potential for usage in evolutionary and conservation studies, we strongly urge against sole use of mtDNA markers for identification of species and hybrids. As the results of this study, as well as that of Malukiewicz et al. (2021)^7^ clearly show, cryptic hybrids can easily be mistaken for species, and had we only depended on mtDNA results we would have misidentified three cryptic *Callithrix* hybrids as *C. jacchus* and *C. penicillata*. Instances of cryptic hybrids have also been shown among natural *C. jacchus* x *C. penicillata* hybrids^24^. All of these instances underline the need to use several lines of evidence for taxanomic identification of marmoset individuals, particularly due to widespread anthropogenic hybridization among marmosets. We used a combination of phenotypic and mitochondrial data to classify the sampled individual BJT022 as a cryptic hybrid. As mitochondrial DNA is maternally transmitted, it is also not possible to genetically identify the paternal lineage of hybrids without further use of autosomal or Y-chromosome genetic markers. When ever phenotypic data are available, these data should be used jointly with molecular data for identification or classification of a marmoset individual as belonging to a specific species or hybrid type. Indeed, the integrated use of phenotypic and molecular approaches will lead to a better understand the phenomena that involve hybridization processes^29^.

### Conclusion

Brazilian legal instruments that protect *C. aurita* consider hybridization a major threat to the survival of this species^12,8^. In this report, we have uncovered the first known case of cryptic hybridization between *C. aurita* and *C. penicillata*, which may represent a larger trend of genetic introgression from exotic into native marmosets in southeastern Brazil. Our findings are based on the combination of two recent innovations in the field of genomics, that of genomic skimming and portable long-read sequencing on the ONT minION. Given that *C. aurita* is still very deficient for genetic data, our approach provides a substantial advance in making more genomic data available for one of the world’s most endangered primates. Genomic skimming based on ONT sequencing can be integrated easily with phenotypic and other genetic data to quickly make new information accessible on species biodiversity and hybridization. Such data can then be utilized within the legal Brazilian framework to protect endangered species like *C. aurita*. More specifically, rapid access to emerging biological information on such species leads to more informed decisions on updating or modifying legal actions for protecting endangered fauna. The ONT genomic skimming approach we present here can be further utilized and optimized to more rapidly generate genomic information without the need for specialized technological infrastructure nor the need for *a priori* genomic information.

## Methods

### Sampling

A skin biopsy was sampled from a single adult male marmoset with a *Callithrix aurita* phenotype at the Guarulhos Municipal Zoo, Guarulhos, Sao Paulo, Brazil following the procedure described in Malukiewicz et al. (2017, 2021)^7, 16^. Tissues were collected under the approval of the ASU Institutional Animal Care and Use Committee Animals (ASU IACUC, protocols #11-1150R, 15-144R) and Brazilian Environmental Ministry (SISBIO protocols #47964-2 and #28075-2). Biological tissue sampling complied with all institutional, national, and international guidelines. The sample is registered in the Brazilian SISGEN database under number A18C1CE.

### Sequencing

The mitochondrial genome of the sampled marmoset was reconstructed using a previously published approach based on long range polymerase chain reaction (L-PCR) and Sanger sequencing following Malukiewicz et al. (2017)^16^ and Malukiewicz et al. (2021)^7^. Then the Oxford Nanopore minION portable sequencer was used in-house for sequencing of the same sample on two R9.4 (Oxford Nanopore FLO-MIN106) flow cells. DNA was prepared following manufacturer instructions for the Ligation Sequencing Kit 1D (Oxford Nanopore SQK-LSK108). Briefly, for each minION sequencing run, 1000 ng of input genomic DNA was used, which was quantified prior with a Qubit 3.0 dsDNA BR assay. The genomic DNA was adjusted to a volume of 46 ul with nuclease-free water (NFW) and then transferred to a Covaris g-Tube for fragmentation into 8 kilobase (kb) fragments. The g-Tube was centrifuged in a minicentrifuge for 1 minute, inverted, and then spun again for 1 minute. One ul of the fragmented DNA was removed to check the fragment size via gel electrophoresis and quantification with the Qubit 3 dsDNA BR assay. End repair and dA-tailing of the fragmented DNA was done with the NEBNext Ultra II End Repair/dA-Tailing Module (NEB E7546S). A reaction mix was made by adding 7 ul of NEB Ultra II end-prep reaction buffer, 3 ul of NEB Ultra II enzyme mix, and 5 ul of NFW to 45 ul of fragmented DNA, and the reaction was incubated at 20°C for 5 min and 65°C for 5 min. The reaction was cleaned up with a 1× volume (60 ul) of AMPure XP beads, and the cleaned DNA was eluted in 31 ul NFW. A 1 ul aliquot was quantified via the Qubit 3 dsDNA BR assay to check that at least 700 ng of DNA were retained. Adapter ligatation was done by mixing 30 ul of end-prepped DNA with 20 ul SQK-LSK108 adapter Mix, and 50 ul NEB Blunt/TA Master Mix (NEB M0367S). The mix was flicked and then incubated at room temperature for 10 min. The adapter-ligated DNA was cleaned up by adding a 0.4× volume (40 ul) of AMPure XP beads, followed by resuspension of the bead pellet in 140 ul ABB buffer (from SQK-LSK108). The beads were then resuspended in 15 ul SQK-LSK108 ELB buffer, incubated at room temperature for 10 min, pelleted again, and then the eluate was transferred to a new tube. One ul of the eluate was quantified by Qubit to check that at least 430 ng of DNA were retained.

Each MinION R9.4 flow cells was primed prior sequencing by loading 800 ul of priming buffer (48% v/v SQK-LSK108 RBF buffer in NFW) into the flow cell priming port, waiting 5 minutes, and then lifting the SpotON sample port cover. Then another 200 ul of priming mix were loaded into the flow cell priming port. Sequencing libraries for both runs were prepared by adding 35 ul RBF buffer, 25.5 ul of Library Loading Beads from the Oxford Nanopore Libary Loading Bead kit (EXP-LLB001), and 2.5 ul NFW to 12 ul of the presequencing mix. A volume of 75 ul of library was loaded drop by drop onto the SpotON sample port and capillary action pulled the library into the flow cell. MinKNOW software 2.2 was used to set up two 48 hours MinION sequencing runs on a MacBook Pro laptop computer and read base calling of resulting reads in FAST5 format and conversion to FASTQ format was done with Guppy 2.1.2. Guppy also generated a series of quality control files which were used to assess the performance of both sequencing runs. Both MinKNOW and Guppy are only available to ONT customers via their community site (https://community.nanoporetech.com).

### ONT Sequencing Data Analysis and Mitogenome Reconstruction and Annotation

To first filter FASTQ reads output by Guppy, we first used PORECHOP 0.2.4^30^ to remove sequencing adapters and with NANOFILT 2.7.1^31^ remove low quality reads (Q<7). Then we identified minION reads representing the mitogenome with the BLAST-based mtBlaster script^5^, using the mitogenome of *C. aurita* (GenBank Accession MN787075) as the reference genome. MtBlaster output was assembled *de novo* into a full mitogenome with the FLYE 2.8.2^32, 33^ assembler and polisher for single molecule sequencing reads such as those produced by the minION. The command line script for the above steps is available at https://github.com/Callithrix-omics/aur_nano.

The assembled minION mitogenome was aligned to the Sanger sequencing results with MAFFT 7.475 (www.ebi.ac.uk/Tools/msa/mafft) for comparison of the two mitogenomes. The alignment was visually inspected in MESQUITE^34^, and merged manually for a final consensus sequence. The reconstructed mitogenome sequence was annotated using the CHLOROBOX web server^18^ with the following parameters: “circular sequence” option, mitochondrial DNA, tRNAscan-SE v2.0 in “Mammalia Mitochondrial tRNAs” mode was enabled, and Server References from NCBI were selected including all RefSeqs for *Callithrix*.

### Mitogenome Phylogenetic and Diversity Analyses

To determine the lineage of the reconstructed mitogenome, we combined this sequence with other *Callithrix* mitogenomes^7^, and used other New World primate GenBank sequences as outgroups. Accession numbers of all sequences used are listed in Table S1. We kept mitochondrial genomes in their entirety, but trimmed part of tRNA-Phe, 12s rRNA and the control region to accommodate the length of all utilized sequences. All mitogenomes were aligned in MAAFT with default settings. A maximum likelihood phylogeny of all sequences was generated with IQTREE 1.6.12^35^. We used the optimal substitution model (TPM2u+F+I+G4) as calculated automatically with ModelFinder^36^ in IQ-TREE under the Bayesian Information Criterion (BIC). We performed the ML analysis in IQ-TREE with 1,000 ultrafast bootstrap (BS) replications^37^. DNASp 6.12.03^38^ was used for population genomic analysis to calculate standard diversity indices.

## Acknowledgements

This work was supported by a Brazilian CNPq Jovens Talentos Postdoctoral Fellowship (302044/2014-0), a Brazilian CNPq DCR grant (300264/2018-6), an American Society of Primatologists Conservation Small Grant, a Marie-Curie Individual Fellowship (AMD-793641-4) and an International Primatological Society Research Grant for JM. This work was also supported by an Arizona State University SOLS/OKED Research Investment grant to RAC, AV, MAW, and JM, as well as an Arizona State University RTI Postdoctoral Grant to JM. Additional internal funding from the German Primate Center to LW and JM also supported this work. The funding agencies had no roles in study design, data collection and analysis, decision to publish, or preparation of the manuscript. We would like to thank the Golden Lion Tamarin Association, CEMAFAUNA, the Guarulhos Municipal Zoo, the Beagle Lab at UFV, and many biologists, field technicians, veterinarians, and other individuals that made this research possible.

## Author contributions statement

JM formulated the idea for the study, collected samples, obtained funding, conducted wet and dry laboratory work, and wrote the original manuscript. RAC, AV, and MAW provided study guidance, obtained funding, and provided logistical support. JAD provided study guidance, and logistical support. CSI gave access and provided logistical support to collect samples from animals kept at Guarulhos Zoo. PAN provided study guidance, and logistical support. LCMP provided study guidance, and logistical support. CRRM and DZ were major contributors in writing the manuscript. DLS performed DNA extractions, methodological optimization, and carried out PCR. ACS was a major contributor in writing the manuscript, provided study guidance, and logistical support. LW and CR gave significant guidance and input in the development of this study. All authors read and approved the final manuscript.

## Supplemental Material

**Figure S1.**
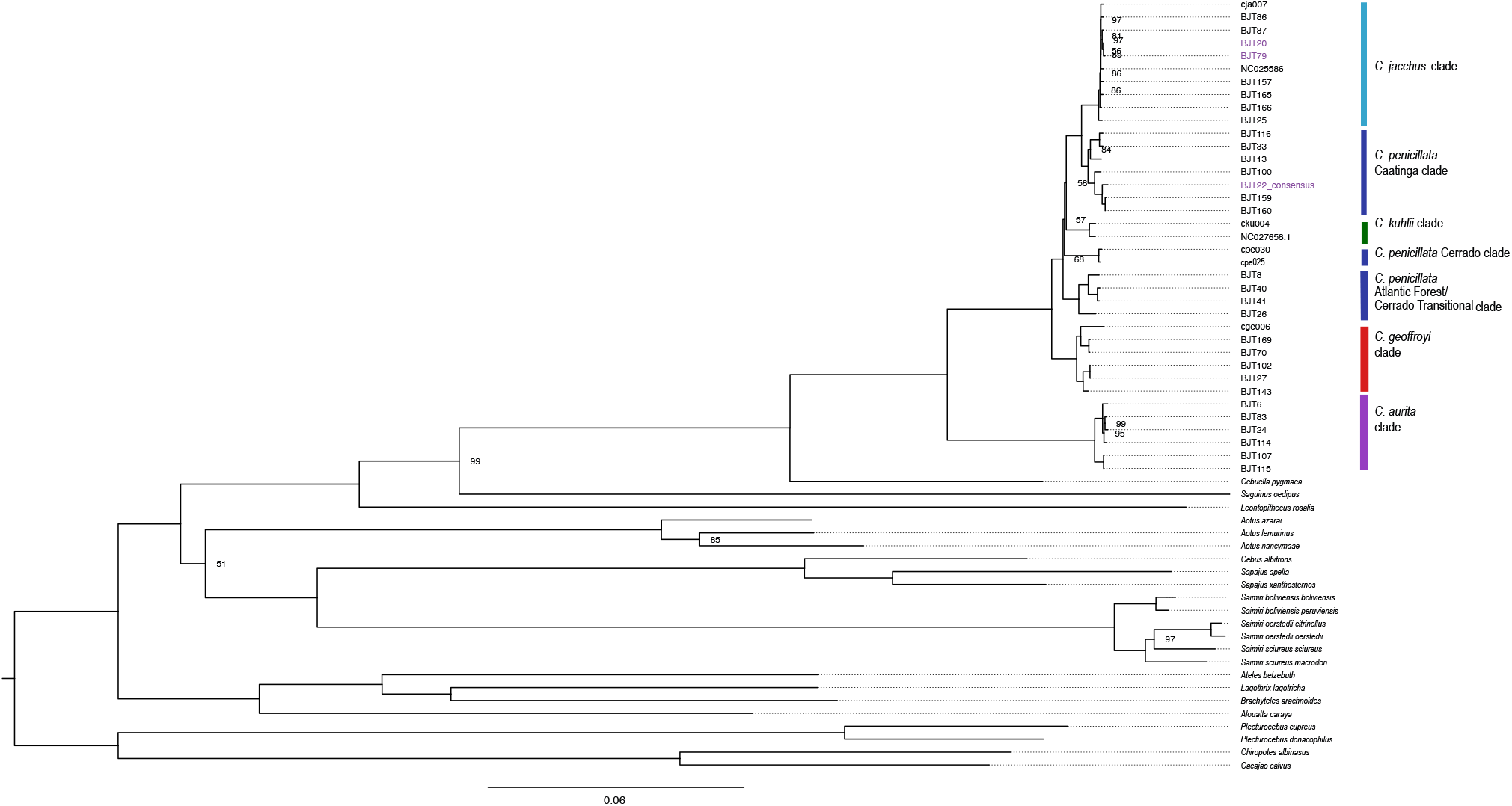
ML tree showing phylogenetic clustering of the *Callithrix* mitogenome of marmoset BJT022 within the *C. penicillata* Caatinga clade. The BJT022 mitogenome is highlighted at the tree tips in purple along with two other mitogenomes of marmosets with *C. aurita* phenotypes from Malukiewicz et al. (2021)^7^ that were incongruent with their mitogenmic lineages. Categorization of major *Callithrix* mitogenomic clades follow that of Malukiewicz et al. (2021)^7^.

**Table S1.**
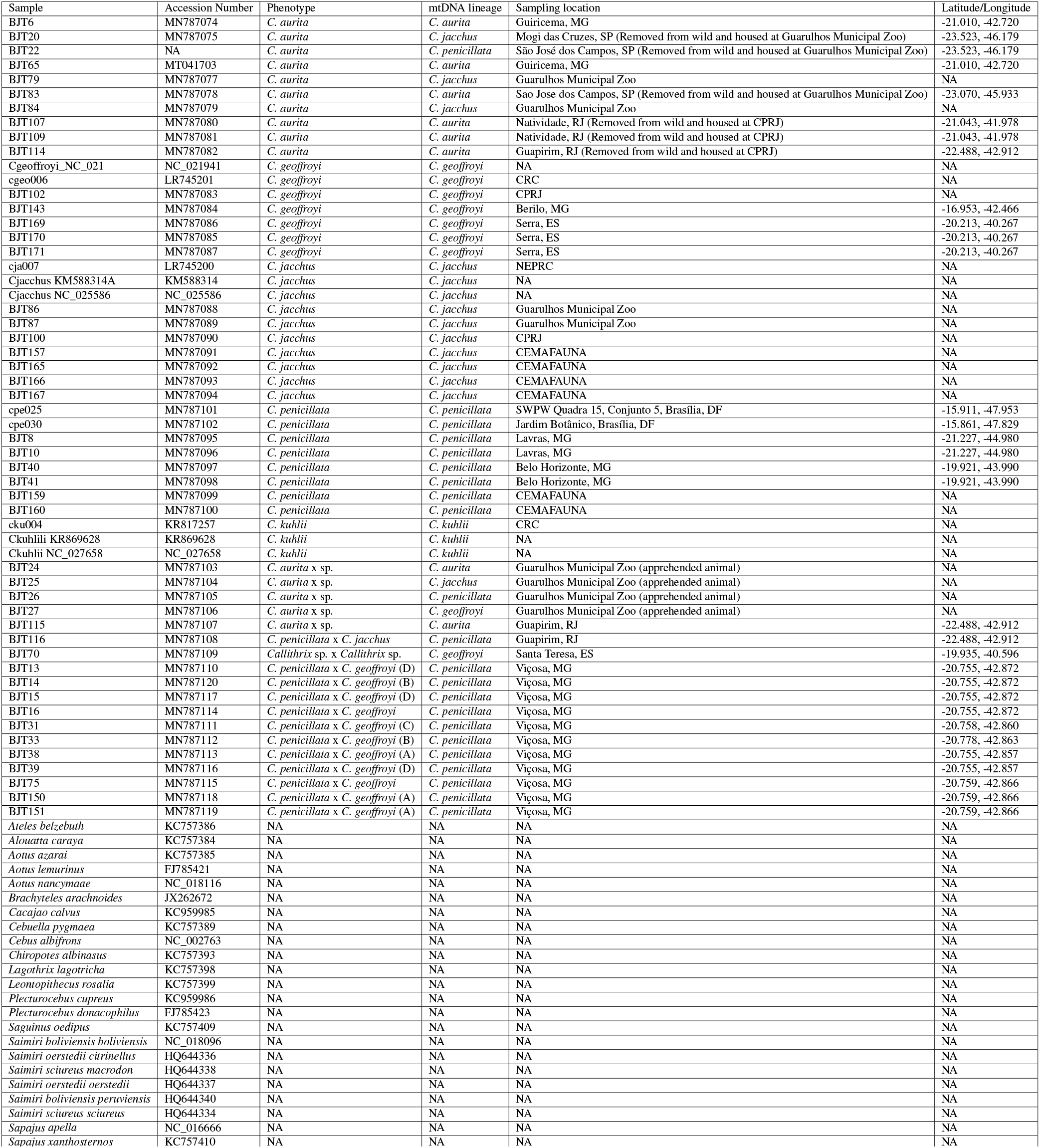
Sample metadata for newly generated *Callithrix* mitogenome from individual BTJ022 and mitogenomic sequences used from Malukiewiczet al. (2021)^7^. The ‘Sample’ column gives ID of each new sampled individual or the species for sequences obtained from previous studies. The ‘Accession’ column gives Genbank accession numbers for each sequence. The ‘Phenotype’ column indicates whether the sampled individual possessed a pure species or hybrid phenotype, and capital letters in parentheses next to *Callithrix penicillata* x *Callithrix geoffroyi* category are specific phenotype classifications following Figure 5 in Fuzessy et al. (2014)^39^. The ‘mtDNA Lineage’ column indicates phylogenetic classification of the mitochondrial genome of the sampled individual. The ‘Sampling Location’ column indicates where each individual was sampled. Nearest cities are located for individuals sampled from the wild, and facilities are indicated for individuals sampled in captivity. The Guarulhos Municipal Zoo is located in Guarulhos, São Paulo, Brazil; CRC (Callitrichid Research Center) is located in Omaha, Nebraska, US; NEPRC (New England Primate Research Center, no longer in operation) was located in Southborough, Massachusetts, US; CPRJ (Centro de Primatologia do Rio de Janeiro) is located in Guapimirim, Rio de Janeiro, Brazil; CEMAFAUNA (Centro de Concervação e Manejo de Fauna da Caatinga) is located in Petrolina, Pernambuco. Abbreviations for Brazilian states in the ‘Sampling Location’ column are as follows: Espírito Santo (ES), Minas Gerais (MG), Rio de Janeiro (RJ), São Paulo (SP). DF is the Brazilian Federal District. NA = No data Available.

